# Prophylactic efficacy of 5-HT_4_R agonists against stress

**DOI:** 10.1101/712786

**Authors:** Briana K. Chen, Indira Mendez-David, Charlene Faye, Alain M. Gardier, Denis J. David, Christine A. Denny

## Abstract

Enhancing stress resilience could protect against stress-induced psychiatric disorders in at-risk populations. We and others have previously reported that (*R*,*S*)-ketamine acts as a resilience enhancing drug (e.g., prophylactic) against stress when administered 1 week *before* stress. While we have shown that the selective 5-hydroxytryptamine (5-HT) (serotonin) reuptake inhibitor (SSRI) fluoxetine (Flx) is ineffective as a prophylactic, it remains to be determined if other serotonergic drugs could be effective prophylactics. Here, we hypothesized that serotonin 4 receptor (5-HT_4_R) agonists could be prophylactic against fear, depressive-like, and/or anxiety-like behavior. We tested if three 5-HT_4_R agonists with varying affinity (e.g., partial or selective agonists) could protect against stress in two mouse strains by utilizing a chronic corticosterone (CORT) administration or a contextual fear conditioning (CFC) paradigm. Mice were administered RS-67,333, prucalopride, or PF-04995274 at varying doses and then 1 week later were subjected to chronic CORT or CFC. Chronic administration of RS-67,333, but not Flx was efficacious as a prophylactic against CORT in C57BL/6NTac mice. A single injection of RS-67,333 attenuated learned fear in male, but not female 129S6/SvEv mice. RS-67,333 was ineffective against stress-induced depressive-like behavior in the forced swim test (FST). A single injection of either prucalopride or PF-04995274 attenuated learned fear and decreased stress-induced depressive-like behavior. These data show that in addition to (*R*,*S*)-ketamine, 5-HT_4_R agonists are also effective prophylactics against stress, suggesting that the 5-HT_4_R may be a novel target for prophylactic drug development.

## INTRODUCTION

Stress exposure is a significant factor for the development of major depressive disorder (MDD) and post-traumatic stress disorder (PTSD). According to the National Comorbidity Study, approximately 60% of men and 51% of women have been exposed to one or more traumatic events during their lifetime. It is estimated that 7.8% of the overall population experiences PTSD at some point in their lives, with females (10.4%) experiencing the disorder at significantly higher rates than males (5.0%) [1]. Traditionally, affective disorders have been treated from a symptom-suppression approach. Existing drugs aim to mitigate the impact of these chronic diseases, but do not cure or prevent the disease itself. However, if drugs were developed that enhance stress resilience, they could potentially be used in at-risk populations to protect against stress-induced psychiatric disorders.

We and others have recently reported that (*R*,*S*)-ketamine acts as a resilience enhancing drug (e.g., prophylactic) against stress when administered 1 week *before* stress in mice [2–6]. In addition, limited data in human patients have demonstrated (*R,S*)-ketamine’s potential in preventing psychiatric disorders such as PTSD [7] and, potentially in a dose-specific manner, post-partum depression (PPD) [8,9]. Prophylactic drug efficacy has been limited to (*R*,*S*)-ketamine until recently when Gould and colleagues reported that group II metabotropic glutamate receptor (mGlu_2/3_) antagonists are also protective [10]. We have previously reported that the SSRI Flx is ineffective as a prophylactic, but it remains to be determined if other serotonergic drugs could be effective prophylactics and/or if the serotonergic system is involved in prophylactic efficacy.

Numerous studies have implicated the 5-HT_4_Rs as a target for treating depression and anxiety. 5-HT_4_Rs are metabotropic G-protein coupled receptors that stimulate the G_s_/cyclic adenosine monophosphate (cAMP)/protein kinase A (PKA) signaling pathway in response to 5-HT [11–14]. 5-HT_4_Rs are highly expressed in the periphery, including the heart and adrenal gland, as well as in the brain in areas such as the amygdala, medial prefrontal cortex (mPFC), nucleus accumbens (NAc), and hippocampus (HPC) [15,16]. 5-HT_4_R knockout mice display increased anxiety-like behavior and depressive-like behavior, while activation of 5-HT_4_Rs stimulates neurogenesis in the HPC and produces rapid-acting antidepressant-like effects [17–20]. However, if and how 5-HT_4_Rs are involved in stress resilience has yet to be determined.

Here, we hypothesized that since 5-HT_4_Rs have been heavily implicated in depression and anxiety, they may have a role in stress resilience. We focused our studies on three 5-HT_4_R agonists with varying affinity. First, RS-67,333 (1-(4-amino-5-chloro-2-methoxyphenyl)-3-[1(n-butyl)-4-piperidinyl]-1-propanone HCl) is a high-affinity 5-HT_4_R partial agonist [21]. This drug is effective in improving behavioral deficits, decreasing the number of amyloid plaques as well as level of Aβ species, and decreasing hippocampal astrogliosis and microgliosis in the 5XFAD mouse model of Alzheimer’s disease (AD) [22]. Second, prucalopride (4-amino-5-chloro-2,3-dihydro-*N*-[1-3-methoxypropyl)-4-piperidinyl]-7-benzofuran carboxamide monohydrochloride) is a selective, high affinity 5-HT_4_R agonist [23]. In 2018, it was approved by the FDA for chronic constipation and is currently being tested for chronic intestinal pseudo-obstruction. Prucalopride has also been tested in two separate clinical trials to investigate its effects on emotional processing in health volunteers after an acute (e.g. single dose) or chronic (e.g. 1 week) administration [24,25]. Third, PF-04995274 (4-[4-[4-Tetrahydrofuran-3-yloxy)-benzo[d]isoxazol-3-yloxymethyl]-piperidin-1-ylmethyl]-tetrahydropyran-4-ol) is a potent, partial 5-HT_4_R agonist [26]. A clinical trial was conducted to evaluate PF-04995274, alone or in combination with donepezil, on scopolamine-induced deficits in psychomotor and cognitive function in healthy adults; however, this trial was terminated, but not due to safety concerns [27]. Currently, a clinical trial is underway to test whether adjunctive administration of PF-04995247 has positive effects on emotional processing and neural activity in mediated, treatment-resistant (TRD) depressed patients compared to placebo [28].

To determine if 5-HT_4_R agonists may be potential prophylactics against stress, we utilized two different stress models (acute and chronic) in two different strains of mice (C57BL/6NTac and 129S6/SvEv). We found that RS-67,333, prucalopride, and PF-04995274 attenuate learned fear. Additionally, prucalopride, and PF-04995274 also decrease stress-induced depressive-like behavior in the FST. These effects were limited to male mice, as we did not determine prophylactic efficacy of RS-67,333 in female 129S6/SvEv mice. These data suggest that the 5-HT_4_R receptor may be a novel target for prophylactic development.

## MATERIALS AND METHODS

### Mice

#### C57BL/6NTac mice

Male C57BL/6NTac mice were purchased from Taconic Farms (Lille Skensved, Denmark) at 8 weeks of age and were housed 5 per cage before the start of CORT treatment. All testing was conducted in compliance with the laboratory animal care guidelines and with protocols approved by the Institutional Animal Care and Use Committee (IACUC) (European Directive, 2010/63/EU for the protection of laboratory animals, permissions # 92-256B, authorization ethical committee CEEA n°26 2012_098). All mice were housed in a 12-h (06:00-18:00) light-dark colony room at 22°C. Food and water were provided *ad libitum*. Behavioral testing was performed during the light phase.

#### 129S6/SvEv mice

Male and female 129S6/SvEvTac mice were purchased from Taconic (Hudson, NY) at 7 weeks of age. The procedures described herein were conducted in accordance with the National Institutes of Health (NIH) regulations and approved by the IACUC of the New York State Psychiatric Institute (NYSPI). All mice were housed in a 12-h (06:00-18:00) light-dark colony room at 22°C. Food and water were provided *ad libitum*. Behavioral testing was performed during the light phase.

#### Drugs

All drugs were prepared in physiological saline and all injections were administered intraperitoneally (i.p.) in volumes of 0.1 cc per 10 mg body weight unless otherwise noted.

### Stress models

#### Corticosterone (CORT) model

In this model, glucocorticoid levels are exogenously increased in C57BL/6NTac mice. This chronic CORT elevation dysregulates the hypothalamic-pituitary-adrenal axis (HPA) in a manner similar to that observed in clinical depression. For example, there is a blunting of the HPA-axis response to stress in CORT-treated mice, as shown by marked attenuation of stress-induced CORT levels [19,29]. This model reliably induces anxious and depressive-like behavior in mice. The dose and duration of CORT treatment was selected based on previous studies [19,29]. CORT (35 μg/ml, equivalent to about 5 mg/kg/day) dissolved in 0.45 % hydroxypropyl-β-cyclodextrin (β-CD) or vehicle (VEH) (0.45% β-CD) was available *ad libitum* in the drinking water in opaque bottles to protect it from light. VEH- and CORT-treated water was changed every 3 days to prevent any possible degradation.

#### Contextual Fear Conditioning (CFC)

A 3-shock CFC procedure was administered as previously published [30,31]. Briefly, mice were placed into context A and administered 3 2-s shocks (0.75 mA) at 180 s, 240 s or 300 s following placement into context A. Mice were removed from the context 15 s following the termination of shock (at 317 s). For context retrieval, mice were placed back into context A for 300 s.

### Drugs

#### Fluoxetine hydrochloride (Flx)

Flx (BioTrend Chemicals AG, BG197) was administered in the drinking water (18 mg/kg/day) for 3 weeks before the start of CORT.

#### RS-67,333 (RS)

RS-67,333 (Tocris, Catalog No. 0989, Minneapolis, MN) was administered chronically or in a single injection. For the chronic experiment, RS-67,333 (1.5 mg/kg/day) was administered via ALZET osmotic minipumps (ALZET, Cupertino, CA). For the acute experiment, RS-67,333 was administered in a single dose of 1.5, 10, or 30 mg/kg of body weight 1 week before the start of CFC. RS-67,333 was dissolved in saline using an ultrasonic homogenizer (BioLogics, Model 3000, Manassas, VA).

#### (*R*,*S*)-ketamine (K)

(*R*,*S*)-ketamine (Ketaset III, Ketamine HCl injection, Fort Dodge Animal Health, Fort Dodge, IA) was administered in a single dose at 30 mg/kg of body weight 1 week before the start of CFC. A dose of 30 mg/kg of body weight was chosen in the 129S6/SvEv experiments, as previous studies indicated that is the effective dose for prophylactic efficacy [3].

#### Prucalopride

Prucalopride (Sigma, Catalog No: 179474-81-8, St. Louis, MO) was administered a single dose at 3 or 10 mg/kg of body weight 1 week before the start of CFC. Prucalopride was dissolved in saline using an ultrasonic homogenizer (BioLogics, Model 3000, Manassas, VA).

#### PF-04995274

PF-04995274 (Sigma, Catalog No. 1331782-27-4, St. Louis, MO) was administered a single dose at 3 or 10 mg/kg of body weight 1 week before the start of CFC. PF-04995274 was dissolved in saline using an ultrasonic homogenizer (BioLogics, Model 3000, Manassas, VA).

#### Osmotic minipump implantation

ALZET osmotic minipumps (Model 2004, 0.25 μl/hr, 28 days) were implanted subcutaneously under isoflurane anesthesia as previously described [30]. Osmotic minipumps were rotated under the skin two to three times per week.

#### Statistical Analysis

Results from data analyses are expressed as means ± SEM. Alpha was set to 0.05 for all analyses. Data were analyzed using GraphPad Prism v7.0. For all experiments, unless otherwise noted, one- or two-way ANOVAs with repeated-measures were applied to the data as appropriate. Significant main effects and/or interactions were followed by Fisher’s PLSD post hoc analysis or unpaired *t*-tests. All main effects, interactions, and p values are listed in **Supplemental Table S01**.

## RESULTS

### Chronic administration of RS-67,333 is prophylactic against stress in male mice

We have previously reported that chronic Flx administration (3 weeks of administration) is not prophylactic against chronic CORT administration [3]. However, it remained to be determined if other serotonergic drugs could act as prophylactics. Here, we administered Flx (18 mg/kg/day) in the drinking water or RS-67,333 (1.5 mg/kg/day) in osmotic minipumps for 3 weeks prior to CORT administration in C57Bl/6NTac male mice followed by a series of behavioral assays, including the elevated plus maze (EPM), novelty-suppressed feeding (NSF), and sucrose splash test (ST) (**Fig. 1a-1b**). CORT increased body weight over the 6-week behavioral protocol, as previously observed [32], (**Fig. 1c-1f**), but this was attenuated by Flx and RS-67,333 administration.

**Fig. 1.**
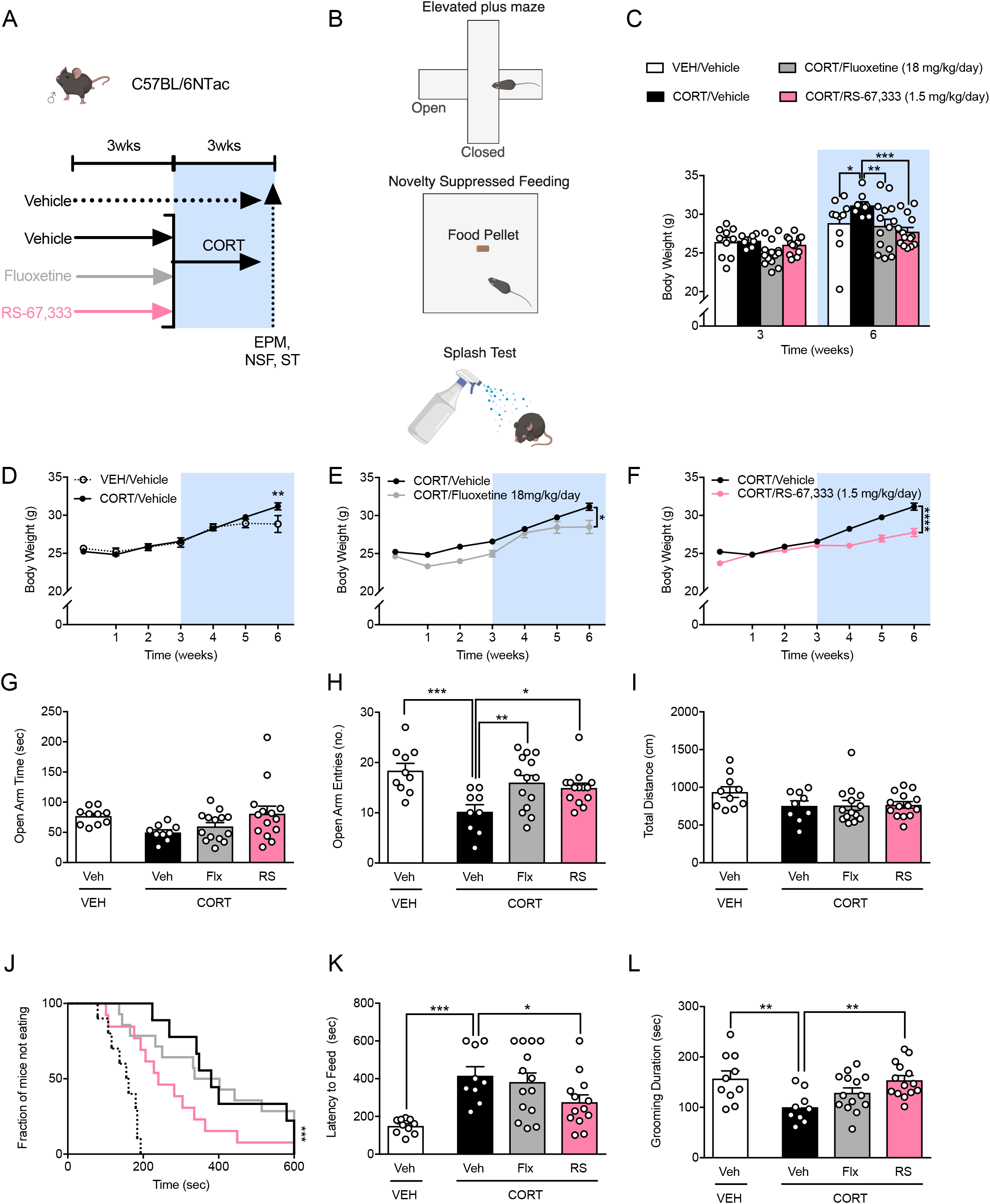
RS-67,333 protects against depressive- and anxiety-like behavior induced with a neuroendocrine model in male C57BL/6NTac mice. (a) Experimental design. Behavioral assays to test anxiety-like behavior (EPM, NSF) and depressive-like behavior (ST). (c-f) By Week 6, CORT administration resulted in increased body weight when compared with VEH administration. RS and Flx administration resulted in decreased body weight in CORT-treated mice. (g) All groups of mice exhibited comparable amounts of time in the open arms of the EPM. (h) CORT + Veh mice had significantly less entries into the open arms of the EPM when compared with VEH + Veh mice. RS, but not Flx administration increased the number of entries into the open arms of the EPM in CORT-treated mice. (i) All groups of mice traveled a similar distance in the EPM. (j-k) CORT administration increased the latency to feed in the NSF when compared with the VEH administration. RS, but not Flx administration decreased the latency to feed in CORT-treated mice. (l) CORT administration decreased grooming duration in the ST when compared with VEH administration. RS, but not Flx administration increased the grooming duration in CORT-treated mice. (n = 9-14 male mice per group). Error bars represent + SEM. *, p < 0.05; ** p < 0.01; ***, p < 0.001; ****, p < 0.0001. VEH, vehicle; Veh, vehicle; CORT, corticosterone; Flx, fluoxetine; RS, RS-67,333; EPM, elevated plus maze; NSF, novelty suppressed feeding; ST, splash test; sec, seconds; no., number; cm, centimeters; g, grams.

In the EPM, CORT + Veh, CORT + Flx, and CORT + RS-67,333 administration did not alter the time spent in the open arms when compared with VEH + Veh administration (**Fig. 1g**). However, CORT + Veh mice exhibited a significantly decreased number of entries into the open arms of the EPM when compared with VEH + Veh mice (**Fig. 1h**). CORT + Flx and CORT + RS-67,333 mice had significantly more entries into the open arms of the EPM when compared with CORT + Veh mice. The total distance traveled in the EPM did not differ between any of the groups (**Fig. 1i**).

Next, the NSF task was administered to assay innate, anxiety-like behavior (**Fig. 1j-1k**). CORT + Veh mice exhibited an increased latency to approach the food pellet when compared with VEH + Veh mice. CORT + RS-67,333, but not CORT + Flx mice exhibited a significantly decreased latency to approach the pellet when compared with CORT + Veh mice.

Finally, in the ST, CORT + Veh mice exhibited decreased grooming duration when compared with VEH + Veh mice (**Fig. 1l**). CORT + RS-67,333, but not CORT + Flx mice exhibited increased grooming duration when compared with CORT + Veh mice. These data suggest that chronic RS-67,333, but not chronic Flx administration is prophylactic against CORT-induced behavioral abnormalities.

### A single injection of RS-67,333 attenuates learned fear and protects against stress-induced hypophagia in male mice

Previously, we have shown that a single injection of (*R*,*S*)-ketamine is prophylactic against stress-induced depressive-like behavior and attenuates learned fear in 129S6/SvEv mice [3]. Here, we sought to determine if a single injection of RS-67,333 could also prevent a variety of maladaptive behaviors following a single, acute stressor. Male 129S6/SvEv mice were injected with saline or RS-67,333 (1.5, 10, or 30 mg/kg) (**Fig. 2a**). One week later, mice were administered a 3-shock CFC paradigm. Mice administered 30, but not 1.5 or 10 mg/kg, of RS-67,333 exhibited significantly less freezing during CFC training when compared with mice administered saline (**Fig. 2b**). Five days later, mice were re-exposed to the training context. Mice administered 1.5 or 10, but not 30 mg/kg of RS-67,333 exhibited significantly less freezing when compared with mice administered saline (**Fig. 2c-2d**).

**Fig. 2.**
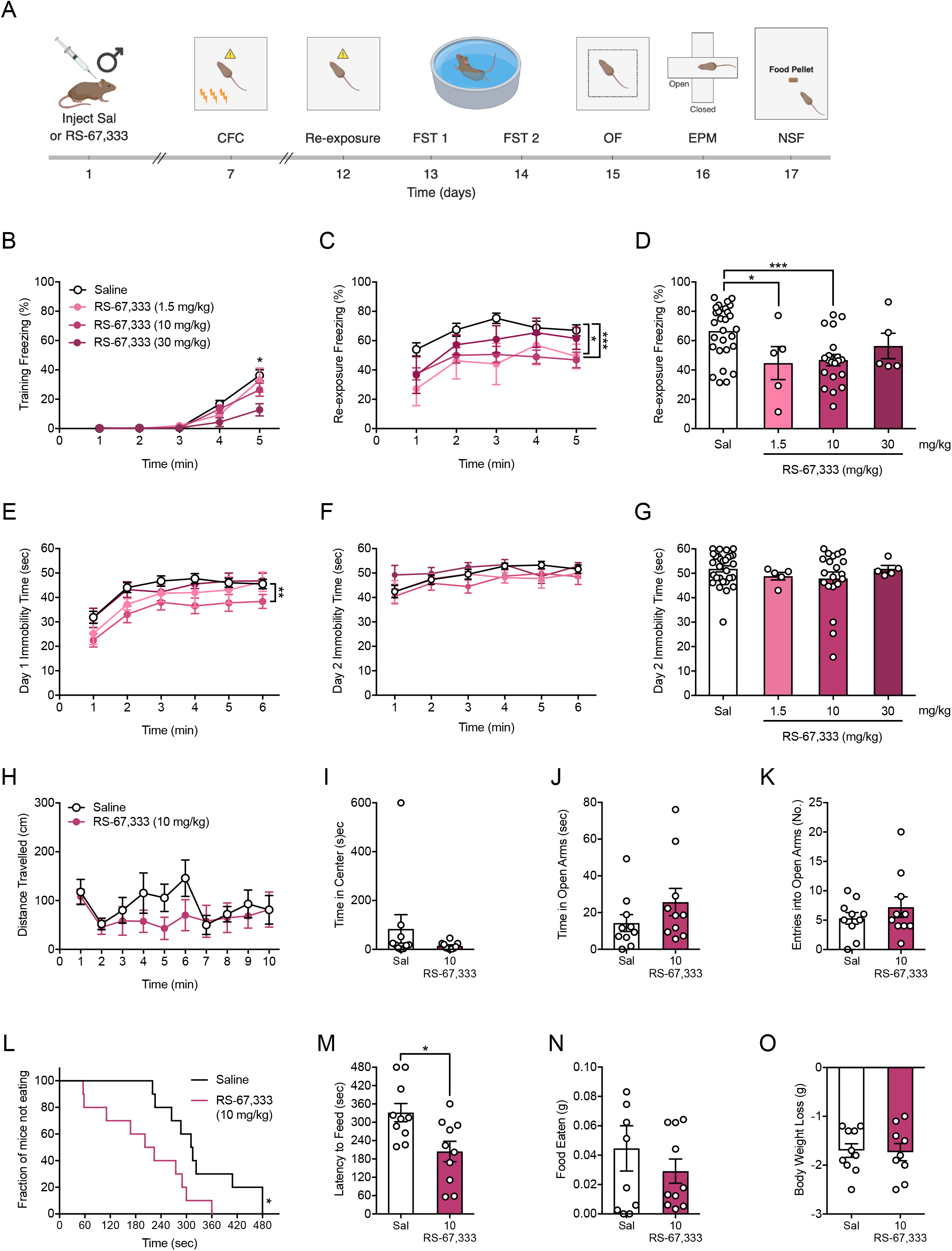
A single, prophylactic injection of RS-67,333 attenuates learned fear and prevents novelty-induced hypophagia in male 129S6/SvEv mice. (a) Experimental design. (b) Mice administered 30, but not 1.5 or 10 mg/kg of RS-67,333 exhibited significantly less freezing during CFC training when compared with mice administered saline. (c-d) Mice administered 1.5 or 10, but not 30 mg/kg of RS-67,333 at exhibited significantly less freezing when compared with mice administered saline. (e) Mice administered 10, but not 1.5 or 30 mg/kg of RS-67,333, exhibited reduced immobility when compared with mice administered saline during day 1 of the FST. (f-g) All groups of mice had comparable amounts of immobility during day 2 of the FST. (h-i) RS-67,333 (10 mg/kg) did not alter distance travelled or time spent in the center of the OF when compared to saline mice. (j-k) Both groups of mice had comparable time spent in the open arms and entries into the open arms of the EPM. (l-m) Mice administered RS-67,333 (10 mg/kg) exhibited a significantly reduced latency to feed when compared to saline mice. (n) Mice in both groups ate a comparable amount of food in the home cage following the NSF. (o) Following food deprivation, mice in both groups lost a comparable amount of weight. (n = 5-29 male mice per group). Error bars represent ± SEM. *, p < 0.05; ***, p < 0.001. Sal, saline; CFC, contextual fear conditioning; FST, forced swim test; min, minutes; sec, seconds; g, grams.

Following CFC, mice were administered the FST. On Day 1, mice administered 10, but not 1.5 or 30 mg/kg, of RS-67,333 were significantly less immobile when compared with saline mice (**Fig. 2e**). However, on Day 2, immobility time was comparable between all groups (**Fig. 2f-2g**).

Next, mice administered saline or RS-67,333 (10 mg/kg) were tested in the OF. Both groups of mice travelled a comparable distance (**Fig. 2h**) and spent a comparable amount of time in the center of the arena (**Fig. 2i**). Subsequently, mice were tested in the EPM, and neither in the open arms nor entries into the open arms of the maze was significantly different between saline or RS-67,333 mice (**Fig. 2j-2k**).

Finally, mice were administered the NSF. Mice given prophylactic RS-67,333 (10 mg/kg) exhibited a significantly reduced latency to approach the pellet (**Fig. 2l-2m**). However, neither food eaten in the home cage nor weight loss following food deprivation differed between the groups (**Fig. 2n-2o**). Together, these data indicate that a single injection of RS-67,333 is effective as a prophylactic in attenuating learned fear and preventing stress-induced hypophagia, but not depressive-like behavior, as measured by the FST, in male 129S6/SvEv mice.

### A single prophylactic injection of RS-67,333 is not prophylactic against stress in female mice

We next sought to determine if a single injection of RS-67,333 could also be prophylactic in female mice. Female 129S6/SvEv mice were injected with saline or RS-67,333 (1.5 or 10 mg/kg) (**Fig. 3a**). One week later, mice were administered a 3-shock CFC paradigm. All groups of mice exhibited comparable levels of freezing during CFC training (**Fig. 3b**). Five days later, mice were re-exposed to the training context. Again, all groups of mice exhibited comparable levels of freezing (**Fig. 3c-3d**).

**Fig. 3.**
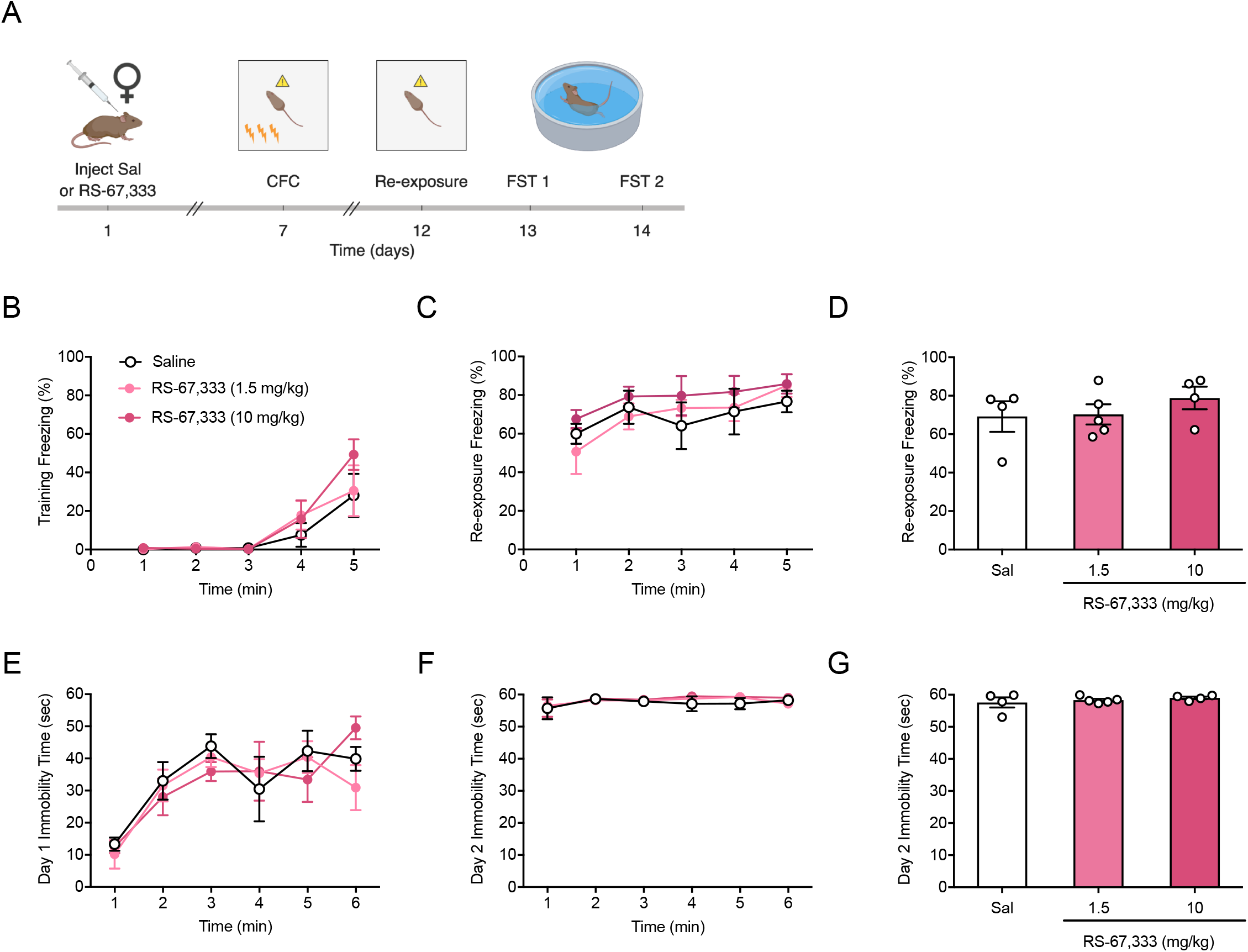
A single, prophylactic administration of RS-67,333 does not attenuate learned fear or decrease depressive-like behavior in female 129S6/SvEv mice. (a) Experimental design. (b) All mice exhibited comparable levels of freezing during CFC training. (c-d) All groups exhibited comparable levels of freezing during re-exposure. (e) All groups of mice had comparable amounts of immobility during day 1 of the FST. (f-g) All groups of mice had comparable amounts of immobility during day 2 of the FST. (n = 4-5 female mice per group). Error bars represent ± SEM.. Sal, saline; CFC, contextual fear conditioning; FST, forced swim test; min, minutes; sec, seconds.

Following CFC, mice were administered the FST. During days 1 (**Fig. 3e**) and 2 (**Fig. 3f-3g**) of the FST, all groups of mice had comparable levels of immobility. These data indicate that a single injection of RS-67,333 is not effective as a prophylactic in attenuating learned fear or in decreasing stress-induced depressive-like behavior in female 129S6/SvEv mice.

### A single prophylactic injection of prucalopride or PF-04995274 is prophylactic against stress in male mice

We next sought to determine if other 5-HT_4_R agonists could also be prophylactic in male 129S6/SvEv mice. Male 129S6/SvEv mice were injected with saline, (*R*,*S*)-ketamine (30 mg/kg), prucalopride (3 or 10 mg/kg), or PF-04995274 (3 or 10 mg/kg) (**Fig. 4a**). One week later, mice were administered a 3-shock CFC paradigm. All groups of mice exhibited comparable levels of freezing during CFC training (**Fig. 4b**). Five days later, mice were re-exposed to the training context. As we have previously published, (*R*,*S*)-ketamine attenuated learned fear (**Fig. 4c-4d**). Interestingly, prucalopride at 3, but not 10 mg/kg and PF04995274 at 10, but not 3 mg/kg, attenuated learned fear when compared with saline administration.

**Fig. 4.**
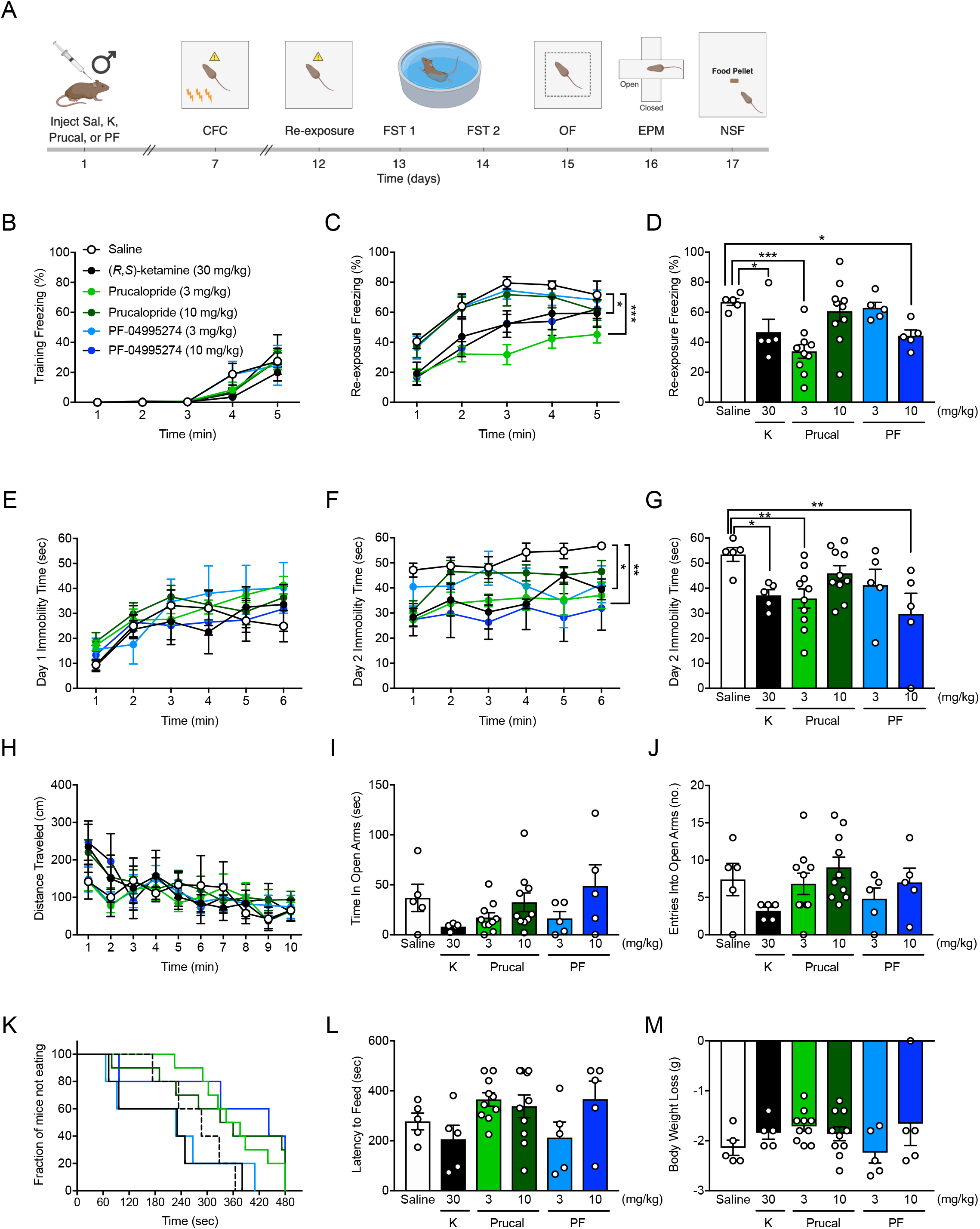
A single, prophylactic administration of prucalopride or PF-04995274 attenuates learned fear and decreases depressive-like behavior in male 129S6/SvEv mice. (a) Experimental design. (b) All mice exhibited comparable levels of freezing during CFC training. (c-d) (*R*,*S*)-ketamine (30 mg/kg), prucalopride (3 mg/kg), and PF-04995274 (10 mg/kg), but not prucalopride (10 mg/kg) or PF-04995274 (3 mg/kg), administration attenuated learned fear when compared with saline administration. (e) All groups of mice had comparable amounts of immobility during day 1 of the FST. (f-g) (*R*,*S*)-ketamine (30 mg/kg), prucalopride (3 mg/kg), and PF-04995274 (10 mg/kg), but not prucalopride (10 mg/kg) or PF-04995274 (3 mg/kg) significantly decreased immobility time during day 2 of the FST. (h) All groups of mice traveled a comparable amount of distance in the OF. (i) All groups of mice spent a comparable amount of time in the open arms of the EPM. (j) All groups of mice had a comparable number of entries into the open arms of the EPM. (k-l) All groups of mice had a comparable latency to approach the pellet in the NSF. (m) All groups of mice lost a comparable amount of weight following food deprivation for the NSF. (n = 5-10 male mice per group). Error bars represent ± SEM. *, p < 0.05; ** p < 0.01; ***, p < 0.001. Sal, saline; K, (*R*,*S*)-ketamine; Prucal, prucalopride; PF, PF-04995274; CFC, contextual fear conditioning; FST, forced swim test; OF, open field; EPM; elevated plus maze; NSF, novelty suppressed feeding; min, minutes; sec, seconds; cm: centimeters; no: number; mg, milligram; kg, kilogram.

Following CFC, mice were administered the FST. During day 1, all groups of mice had comparable levels of immobility (**Fig. 4e**). During day 2, (*R*,*S*)-ketamine administration decreased immobility time when compared with saline administration (**Fig. 4f-4g**). Moreover, prucalopride at 3, but not 10 mg/kg and PF04995274 at 10, but not 3 mg/kg decreased immobility time when compared with saline administration.

Stress-induced anxiety-like behavior was next quantified. In the OF, all groups of mice traveled a comparable distance (**Fig. 4h**). In the EPM, all groups of mice spent comparable time in the open arms (**Fig. 4i**) and entered into the open arms a comparable number of times (**Fig. 4j**). In the NSF paradigm, all groups of mice approached the pellet in a comparable amount of time (**Fig. 4k-4l**). Finally, all mice lost a comparable amount of weight during the NSF paradigm, suggesting that prucalopride and PF04995274 do not impact weight loss (**Fig. 4m**). In summary, these data indicate that a single injection of prucalopride or PF0499574, two 5-HT_4_R agonists, results in prophylactic efficacy by attenuating learned fear and decreasing stress-induced depressive-like behavior. However, these drugs are not prophylactic against stress-induced anxiety-like behavior.

## DISCUSSION

Here, we hypothesized that novel 5-HT_4_R agonists could be prophylactic against fear, depressive-like, and/or anxiety-like behavior. We tested if three 5-HT_4_R agonists with varying affinity (e.g., partial or selective agonists) could protect against stress in two mouse strains by utilizing a chronic CORT administration or a CFC paradigm. Chronic administration of RS-67,333 was efficacious as a prophylactic against CORT stress. A single injection of RS-67,333 attenuated learned fear and prevented stress-induced hypophagia in the NSF in male, but not female 129S6/SvEv mice. RS-67,333 was ineffective against stress-induced depressive-like behavior, as measured by the FST, and anxiety-like behavior. A single injection of either prucalopride or PF-04995274 attenuated learned fear and decreased depressive-like behavior but had no effect on anxiety-like behavior. These data show that in addition to (*R*,*S*)-ketamine, 5-HT_4_R agonists are also effective prophylactics against stress, suggesting that the 5-HT_4_R receptor may be a novel target for prophylactic drug development.

The three 5-HT_4_R agonists chosen in this study have differential affinity to the 5-HT_4_R (**Fig. 5**). RS-67,333 and PF-04995274 are high affinity 5-HT_4_R partial agonists, whereas prucalopride is a selective, high affinity 5-HT_4_R agonist. RS-67,333 attenuated learned fear and protected against novelty-induced hypophagia, but did not decrease stress-induced depressive-like behavior. Prucalopride and PF-04995274 attenuated learned fear and decreased depressive-like behavior but had no effect on various measures of anxiety-like behavior. These data suggest that the unique combination of high pK_i_ and partial selectivity for the 5-HT_4_R exhibited by RS-67,333 is sufficient to prevent against anxiety-like behavior whereas the differential activity of prucalopride and PF-04995274 at the 5-HT_4_R protect against stress-induced depressive-like behavior. Further study is necessary to determine if and how the 5-HT_4_R may contribute to the neurobiological mechanisms underlying stress resilience.

**Fig. 5.**
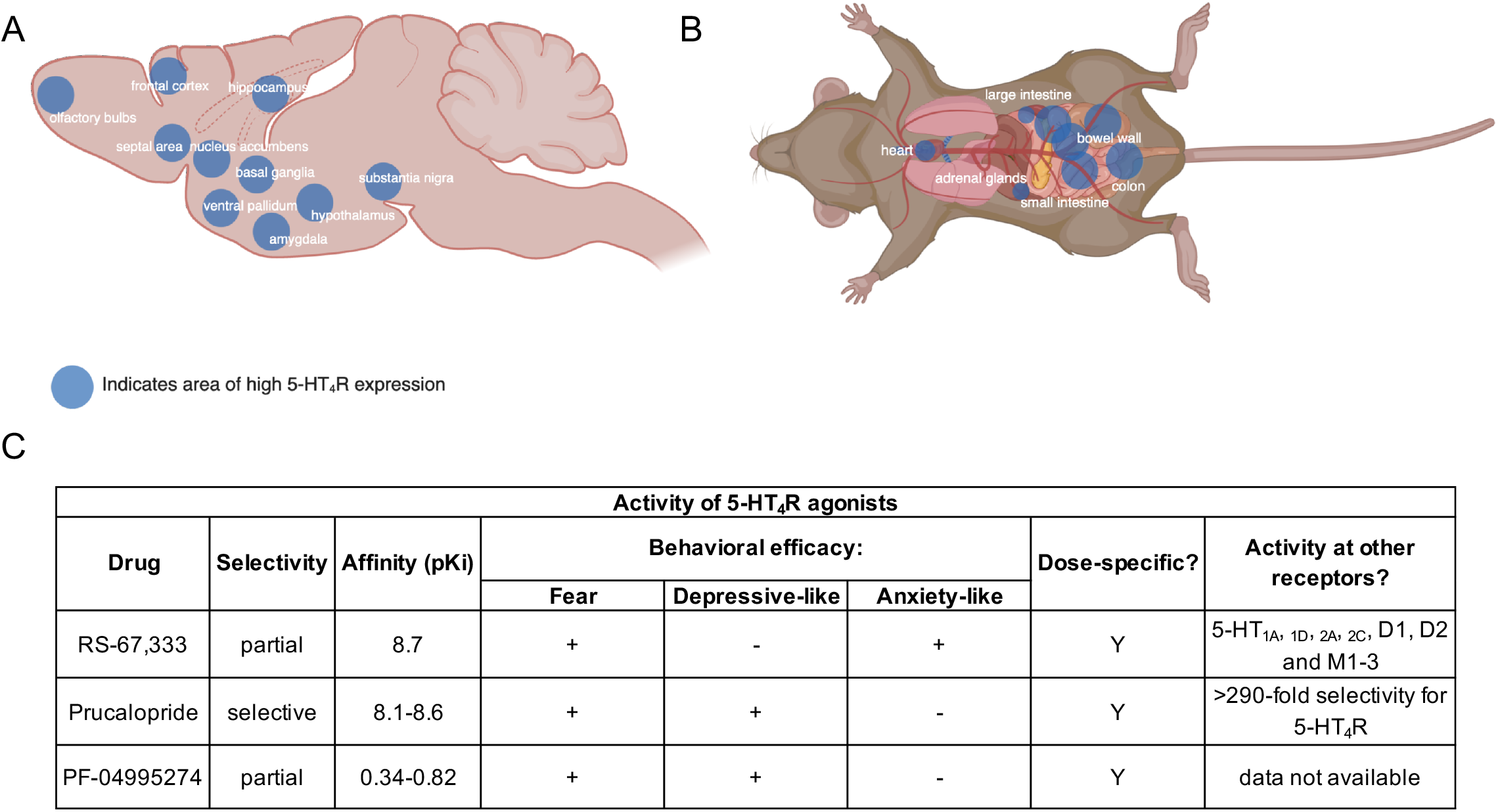
5-HT_4_R expression and summary of behavioral results. (a) The 5-HT_4_R is widely distributed throughout the brain and heavily expressed in areas related to emotional regulation and cognitive function. (b) The 5-HT_4_R is also heavily expressed throughout the periphery and plays a crucial role in regulating ENS activity and function. Summary of behavioral results. RS-67,333, Prucalopride, and PF-04995274 have varying selectivity and affinity for the 5-HT_4_R. These differences may contribute to the drugs’ prophylactic efficacy in preventing fear, depressive-like, or anxiety-like behavior following stress.

The expression and activity of 5-HT_4_Rs within the central nervous system (CNS) and periphery may provide insight into these mechanisms. In the brain, 5-HT_4_Rs are highly expressed in areas of the brain heavily involved in processing emotion, including the HPC, AMG, and PFC [11,15,20,33,34]. In addition to a multitude of other functions, such as modulating dopamine and acetylcholine release [34] as well as facilitating synaptic plasticity [34], 5-HT_4_Rs are known to interact with the calcium effector protein p11 [35]. More specifically, 5-HT_4_Rs are highly co-expressed with p11, which increases surface expression of the receptor in the HPC and AMG, facilitates its downstream signaling pathways, and is necessary for the antidepressant effects of 5-HT_4_R stimulation [35,36]. Moreover, levels of p11 are significantly correlated with measures of suicidality and PTSD, indicating its potential as a biomarker for current suicidal ideation (CSI) and PTSD [37–39]. Additionally, 5-HT4R expression and activity in the PFC is regulated by casein kinase 2 (CK2), which may be an important modulator of depressive- and anxiety-like behaviors [40]. Further studies examining 5-HT_4_R agonists and their effects on these cellular regulators of 5-HT_4_R expression and activity could yield further insight into the protective properties of RS-67,333, prucalopride, and PF-04995274.

In addition to the actions of 5-HT_4_R agonists within the brain, it is likely that these compounds exert additional changes within the periphery. 5-HT_4_Rs are expressed in a wide variety of peripheral areas of the body, such as the enteric nervous system (ENS), adrenal glands, and heart [16]. Importantly, 5-HT_4_Rs play a major role in maintaining communication along the gut-brain axis. Recent data indicate that microbiota in the ENS communicate with the central nervous system (CNS) by stimulating 5-HT_4_Rs present throughout the gut to stimulate serotonin release in the brain [41]. Concurrently, numerous previous studies have shown that activation of 5-HT_4_Rs is neuroprotective against oxidative stress, reduces inflammation, and stimulates neurogenesis in the brain and ENS [41–43]. Our manipulations may have stimulated gut-brain communication to promote neuroprotection and neurogenesis and thereby, enhance resilience against stress-induced fear, depressive-like, and/or anxiety-like behavior. We hypothesize that this action may have had an additive effect on the numerous, well-characterized consequences of 5-HT_4_R stimulation in the brain, such as increasing neuronal firing in the medial PFC (mPFC) and enhancing mitogenesis in the HPC [18,33], although this remains to be determined in future studies.

To develop safe and efficacious pharmacological methods of enhancing stress resilience it will be necessary to determine the toxicity of 5-HT_4_R agonists. Because 5-HT_4_Rs are so widely expressed throughout the periphery, chronic exposure to these drugs could result in negative outcomes [16]. We found that repeated dosing of RS-67,333 did not result in adverse side effects. However, because we did not conduct additional assays, such as assessing changes in cardiovascular activity or liver toxicity, it is impossible to know if repeated 5-HT_4_R agonist exposure would be harmful to peripheral organs. Nonetheless, the drugs that we tested were efficacious in enhancing stress resilience even after a single dose, indicating that 5-HT_4_Rs may be a critical target for developing prophylactic compounds. Further study will be necessary to determine whether single-dose 5-HT_4_R agonists can be developed to prevent stress-induced anxiety-like behavior.

Utilization of numerous mouse strains is essential in determining drug efficacy. While C57BL/6 mice are the most widely used inbred strain, these mice may not be optimized to model susceptibility to stress. Numerous previous reports have shown opposing conclusions if mice of varying genetic backgrounds are tested in the same biological and behavioral assays [44]. These studies suggest that phenotypic relationships cannot be inferred by studying a single genetic background. Here, we utilized two strains - C57BL/6NTac (e.g., resilient) and 129S6/SvEv (e.g., susceptible) mice – in order to validate our effects of RS,67-333 in both a neuroendocrine model of stress and a fear-based stressor. In the C57BL/6NTac mice, we found prophylactic RS-67,333 was effective at decreasing depressive- and anxiety-like behavior, whereas in the 129S6/SvEv mice, we found prophylactic RS-67,333 was effective at attenuating learned fear, but not decreasing depressive-like behavior. Moving forward, numerous strains should be utilized in stress studies in order to infer drug-behavioral relationships.

Despite numerous studies showing prophylactic efficacy in male rodents, only one study to date has shown prophylactic efficacy in female rodents [45]. Maier and colleagues showed that (*R*,*S*)-ketamine administered 1 week prior to an uncontrollable stressor reduced stress-induced activation of the dorsal raphe nucleus (DRN) and eliminated DRN-dependent juvenile social exploration deficits in female rats. However, this study did not measure fear, depressive-like, and anxiety-like behavior in the rodents as done here. Nonetheless, we show that RS-67,333 is an ineffective prophylactic in female 129S6/SvEv mice at the varying doses administered. Prophylactic efficacy of RS-67,333 was not performed in C57BL/6NTac mice, as a previous study of our own has shown that female C57BL/6NTac mice are insensitive to chronic CORT [46]. Future studies are necessary to determine the sex- and dose-specific effects of prophylactic compounds, as twice as many women as men have a lifetime diagnosis of MDD [47].

Overall, the present study has identified three novel compounds to be effective prophylactics against two types of stress. While prophylactics were not effective in female mice, they were in male mice, in accordance with previous (*R*,*S*)-ketamine studies. These data suggest that the 5-HT_4_R may be a novel target for prophylactic development and future studies may lead to novel insights on how 5-HT_4_R agonists administered prior to a stressor result in stress resiliency.

## ACKNOWLEDGMENTS

BKC was supported by a T32 training grant. IM-D was supported by a National Alliance for Research on Schizophrenia and Depression (NARSAD) 2017 Young Investigator Award from the Brain & Behavior Research Foundation and by the Deniker Foundation. CAD was supported by a NIH DP5 OD017908-01. We thank members of the laboratory for insightful comments on this project and manuscript. Additionally, we thank LV Domergue and the staff of the animal care facility of the SFR-UMS Institut Paris-Saclay Innovation Thérapeutique for their technical support.

