## Supplemental Information for "Prophylactic efficacy of 5-HT_4_R agonists against stress"

### SUPPLEMENTAL MATERIALS AND METHODS

#### Behavior:

**Elevated Plus Maze (EPM):** Testing was performed as previously described [1]. Briefly, the maze is a plus-cross-shaped apparatus consisting of four arms, two open and two enclosed by walls, linked by a central platform at a height of 50 cm from the floor. Mice were individually placed in the center of the maze facing an open arm and were allowed to explore the maze for 5 min. The time spent in and the number of entries into the open arms were used as an anxiety index. Videos were scored using ANY-maze (Stoelting Co., Wood Dale, IL) or EPM3C (Bioseb, Vitrolles, France).

**Novelty Suppressed Feeding (NSF):** Testing was performed as previously described [1]. Briefly, the testing apparatus consisted of a plastic box (50 x 50 x 20 cm). The floor of which was covered with approximately 2 cm of wooden bedding and the arena was brightly lit (approximately 1000 lux). For 129S6/SvEv experiments, mice were food restricted for 12 hours. For the C57BL/6NTac experiments, mice were food restricted for 24 h. All food was removed from the home cage. At the time of testing, a single pellet of food (regular chow) was placed on a white paper platform positioned in the center of the box. Each animal was placed in a corner of the box, and a stopwatch was immediately started. The latency of the mice to begin eating was recorded. Immediately after the latency was recorded, the food pellet was removed from the arena. The mice were then placed into their home cage and the amount of food consumed in 5 min was measured (home cage consumption), followed by an assessment of post-restriction weight. A Kaplan-Meier survival analysis was used due to the lack of normal distribution of data.

The Mantel-Cox log-rank test was used to evaluate differences between the experimental groups.

**Splash Test (ST):** This test consisted of squirting 200 μl of a 10% sucrose solution on the mouse's snout as previously described [2]. The grooming duration was quantified using Stopwatch+ (Center for Behavioral Neuroscience, Georgia State University).

**Forced Swim Test (FST):** The FST is typically used in rodents to screen for potential human antidepressants [3,4]. In the FST, time spent immobile, as opposed to swimming, is used as a measure of depressive-like behavior. The FST was administered as previously described [5]. Briefly, mice were placed into clear plastic buckets 20 cm in diameter and 23 cm deep filled 2/3 of the way with 22°C water. Mice were videotaped from the side for 6 min and were exposed to the swim test on 2 consecutive days. Immobility time was scored by an experimenter blind to the experimental groups.

**Open Field (OF):** The OF was administered as previously described [6] with the exception that videos were analyzed using LimeLight (ActiMetrics, Wilmette, IL). Total distance traveled in each quadrant was quantified in 1-minute bins.

### SUPPLEMENTAL FIGURE LEGENDS

Table S01. Statistical analysis summary.
