## Supplemental Table 1 for "Prophylactic efficacy of 5-HT_4_R agonists against stress"

| Table S01: Statistical analysis | | | | | | | | | | |
|---|---|---|---|---|---|---|---|---|---|---|
| Cohort | Behavioral Paradigm | Abbrev | Measurement | Statistical Test | Comparison | F | ° of freedom | p | * | Fig. |
| Figure 1 Chronic RS67333 | Body Weight | BW | Body Weight Change (g) | RMANOVA | Drug | 3.086 | 3,43 | 0.0371 | * | 1C |
| | | | | | Time | 73.890 | 1,43 | <0.0001 | **** | |
| | | | | | Drug x Time | 3.102 | 3,43 | 0.0364 | * | |
| | | | Week 3 | Fisher's LSD | Vehicle vs. CORT/Vehicle | - | - | 0.8643 | ns | |
| | | | | | Vehicle vs. CORT/Fluoxetine 18mg/kg/day | - | - | 0.1100 | ns | |
| | | | | | Vehicle vs. CORT/RS67333 1.5mg/kg/day | - | - | 0.6933 | ns | |
| | | | | | CORT/Vehicle vs. CORT/Fluoxetine 18mg/kg/day | - | - | 0.0838 | ns | |
| | | | | | CORT/Vehicle vs. CORT/RS67333 1.5mg/kg/day | - | - | 0.5717 | ns | |
| | | | | | CORT/Fluoxetine 18mg/kg/day vs. CORT/RS67333 1.5mg/kg/day | - | - | 0.1852 | ns | |
| | | | Week 6 | Fisher's LSD | Vehicle vs. CORT/Vehicle | - | - | 0.0216 | * | |
| | | | | | Vehicle vs. CORT/Fluoxetine 18mg/kg/day | - | - | 0.6945 | ns | |
| | | | | | Vehicle vs. CORT/RS67333 1.5mg/kg/day | - | - | 0.2218 | ns | |
| | | | | | CORT/Vehicle vs. CORT/Fluoxetine 18mg/kg/day | - | - | 0.0048 | ** | |
| | | | | | CORT/Vehicle vs. CORT/RS67333 1.5mg/kg/day | - | - | 0.0004 | *** | |
| | | | | | CORT/Fluoxetine 18mg/kg/day vs. CORT/RS67333 1.5mg/kg/day | - | - | 0.3620 | ns | |
| | | | VEH/Vehicle vs. CORT/Vehicle | RMANOVA | Drug | 0.485 | 1,17 | 0.4957 | ns | 1D |
| | | | | | Time | 55.620 | 6,102 | <0.0001 | **** | |
| | | | | | Drug x Time | 3.201 | 6,102 | 0.0064 | ** | |
| | | | | Fisher's LSD | Week 1 | - | - | 0.5657 | ns | |
| | | | | | Week 2 | - | - | 0.6143 | ns | |
| | | | | | Week 3 | - | - | 0.8876 | ns | |
| | | | | | Week 4 | - | - | 0.8113 | ns | |
| | | | | | Week 5 | - | - | 0.7919 | ns | |
| | | | | | Week 6 | - | - | 0.2446 | ns | |
| | | | | | Week 7 | - | - | 0.0014 | ** | |
| | | | CORT/Veh vs. CORT/F 18mg/kg/day | RMANOVA | Drug | 5.701 | 1,21 | 0.0264 | * | 1E |
| | | | | | Time | 98.570 | 6,126 | <0.0001 | **** | |
| | | | | | Drug x Time | 2.608 | 6,126 | 0.0204 | * | |
| | | | | Fisher's LSD | Week 1 | - | - | 0.4127 | ns | |
| | | | | | Week 2 | - | - | 0.0437 | * | |
| | | | | | Week 3 | - | - | 0.0114 | * | |
| | | | | | Week 4 | - | - | 0.0310 | * | |
| | | | | | Week 5 | - | - | 0.4880 | ns | |
| | | | | | Week 6 | - | - | 0.0799 | ns | |
| | | | | | Week 7 | - | - | 0.0004 | *** | |
| | | | CORT/Veh vs. CORT/RS 1.5mg/kg/day | RMANOVA | Drug | 11.480 | 1,21 | 0.0028 | ** | 1F |
| | | | | | Time | 174.100 | 6,126 | <0.0001 | **** | |
| | | | | | Drug x Time | 22.420 | 6,126 | <0.0001 | **** | |
| | | | | Fisher's LSD | Week 1 | - | - | 0.0042 | ** | |
| | | | | | Week 2 | - | - | 0.8708 | ns | |
| | | | | | Week 3 | - | - | 0.3732 | ns | |
| | | | | | Week 4 | - | - | 0.3215 | ns | |

| | | | | | | | | | | |
|---|---|---|---|---|---|---|---|---|---|---|
| | | | | | Week 5 | - | - | <0.0001 | **** | |
| | | | | | Week 6 | - | - | <0.0001 | **** | |
| | | | | | Week 7 | - | - | <0.0001 | **** | |
| | Elevated Plus Maze | EPM | Open Arm Time | ANOVA | Drug | 2.554 | 3,43 | 0.0678 | ns | 1G |
| | | | Open Arm Entries (No.) | ANOVA | Drug | 5.548 | 3,43 | 0.0026 | ** | 1H |
| | | | | Fisher's LSD | Vehicle vs. CORT/Vehicle | - | - | 0.0003 | *** | |
| | | | | | Vehicle vs. CORT/Fluoxetine 18mg/kg/day | - | - | 0.2031 | ns | |
| | | | | | Vehicle vs. CORT/RS67333 1.5mg/kg/day | - | - | 0.0684 | ns | |
| | | | | | CORT/Vehicle vs. CORT/Fluoxetine 18mg/kg/day | - | - | 0.0043 | ** | |
| | | | | | CORT/Vehicle vs. CORT/RS67333 1.5mg/kg/day | - | - | 0.0182 | * | |
| | | | | | CORT/Fluoxetine 18mg/kg/day vs. CORT/RS67333 1.5mg/kg/day | - | - | 0.5307 | ns | |
| | | | Total Distance | ANOVA | Drug | 1.840 | 3,43 | 0.1542 | ns | 1I |
| | Novelty Suppressed Feeding | NSF | Fraction of mice eating | Log-rank (Mantel-Cox) test | Drug | - | - | <0.0001 | *** | 1J |
| | | | Latency to Feed (sec) | ANOVA | Drug | 7.528 | 3,43 | 0.0004 | | 1K |
| | | | | Fisher's LSD | Vehicle vs. CORT/Vehicle | | | 0.0002 | *** | |
| | | | | | Vehicle vs. CORT/Fluoxetine 18mg/kg/day | | | 0.0003 | *** | |
| | | | | | Vehicle vs. CORT/RS67333 1.5mg/kg/day | | | 0.0388 | * | |
| | | | | | CORT/Vehicle vs. CORT/Fluoxetine 18mg/kg/day | - | - | 0.5809 | ns | |
| | | | | | CORT/Vehicle vs. CORT/RS67333 1.5mg/kg/day | - | - | 0.0267 | * | |
| | | | | | CORT/Fluoxetine 18mg/kg/day vs. CORT/RS67333 1.5mg/kg/day | - | - | 0.0556 | ns | |
| | Splash Test | ST | Grooming Duration (sec) | ANOVA | Drug | 4.909 | 3,43 | 0.0051 | ** | 1L |
| | | | | Fisher's LSD | Vehicle vs. CORT/Vehicle | - | - | 0.0021 | ** | |
| | | | | | Vehicle vs. CORT/Fluoxetine 18mg/kg/day | - | - | 0.0784 | ns | |
| | | | | | Vehicle vs. CORT/RS67333 1.5mg/kg/day | - | - | 0.8398 | ns | |
| | | | | | CORT/Vehicle vs. | - | - | 0.0847 | ns | |
| | | | | | CORT/Vehicle vs. | - | - | 0.0019 | ** | |
| | | | | | CORT/Fluoxetine 18mg/kg/day vs. CORT/RS67333 1.5mg/kg/day | - | - | 0.0868 | ns | |
| | | | Training Freezing (%) | RMANOVA | Drug | 2.5420 | 3,55 | 0.0656 | ns | |
| | | | | | Time | 45.9700 | 4,220 | <0.0001 | **** | |
| | | | | | Drug x Time | 2.064 | 12,220 | 0.0204 | * | |
| | | | | | Saline vs. RS67333 (1.5 mg/kg) | - | - | 0.9785 | ns | |
| | | | | | Saline vs. RS67333 (10 mg/kg) | - | - | 0.9758 | ns | |

| | | | | | | | | | | |
|---|---|---|---|---|---|---|---|---|---|---|
| Figure 2 Acute RS67333 Male | Contextual Fear Conditioning | CFC | Training Freezing (min 1) | Fisher's LSD | Saline vs. RS67333 (30 mg/kg) | - | - | 0.9378 | ns | 2B |
| | | | | | RS67333 (1.5 mg/kg) vs. RS67333 (10 mg/kg) | - | - | 0.9933 | ns | |
| | | | | | RS67333 (1.5 mg/kg) vs. RS67333 (30 mg/kg) | - | - | 0.9359 | ns | |
| | | | | | RS67333 (10 mg/kg) vs. RS67333 (30 mg/kg) | - | - | 0.9257 | ns | |
| | | | Training Freezing (min 2) | Fisher's LSD | Saline vs. RS67333 (1.5 mg/kg) | - | - | 0.9521 | ns | |
| | | | | | Saline vs. RS67333 (10 mg/kg) | - | - | 0.9351 | ns | |
| | | | | | Saline vs. RS67333 (30 mg/kg) | - | - | 0.9521 | ns | |
| | | | | | RS67333 (1.5 mg/kg) vs. RS67333 (10 mg/kg) | - | - | 0.9160 | ns | |
| | | | | | RS67333 (1.5 mg/kg) vs. RS67333 (30 mg/kg) | - | - | >0.9999 | ns | |
| | | | | | RS67333 (10 mg/kg) vs. RS67333 (30 mg/kg) | - | - | 0.9160 | ns | |
| | | | Training Freezing (min 3) | Fisher's LSD | Saline vs. RS67333 (1.5 mg/kg) | - | - | 0.7649 | ns | |
| | | | | | Saline vs. RS67333 (10 mg/kg) | - | - | 0.9687 | ns | |
| | | | | | Saline vs. RS67333 (30 mg/kg) | - | - | 0.9978 | ns | |
| | | | | | RS67333 (1.5 mg/kg) vs. RS67333 (10 mg/kg) | - | - | 0.7547 | ns | |
| | | | | | RS67333 (1.5 mg/kg) vs. RS67333 (30 mg/kg) | - | - | 0.8205 | ns | |
| | | | | | RS67333 (10 mg/kg) vs. RS67333 (30 mg/kg) | - | - | 0.9797 | ns | |
| | | | Training Freezing (min 4) | Fisher's LSD | Saline vs. RS67333 (1.5 mg/kg) | - | - | 0.1524 | ns | |
| | | | | | Saline vs. RS67333 (10 mg/kg) | - | - | 0.2793 | ns | |
| | | | | | Saline vs. RS67333 (30 mg/kg) | - | - | 0.0165 | * | |
| | | | | | RS67333 (1.5 mg/kg) vs. RS67333 (10 mg/kg) | - | - | 0.4480 | ns | |
| | | | | | RS67333 (1.5 mg/kg) vs. RS67333 (30 mg/kg) | - | - | 0.4553 | ns | |
| | | | | | RS67333 (10 mg/kg) vs. RS67333 (30 mg/kg) | - | - | 0.0892 | ns | |
| | | | Training Freezing (min 5) | Fisher's LSD | Saline vs. RS67333 (1.5 mg/kg) | - | - | 0.5384 | ns | |
| | | | | | Saline vs. RS67333 (10 mg/kg) | - | - | 0.0017 | ** | |
| | | | | | Saline vs. RS67333 (30 mg/kg) | - | - | <0.0001 | **** | |
| | | | | | RS67333 (1.5 mg/kg) vs. RS67333 (10 mg/kg) | - | - | 0.2137 | ns | |
| | | | | | RS67333 (1.5 mg/kg) vs. RS67333 (30 mg/kg) | - | - | 0.0025 | ** | |
| | | | | | RS67333 (10 mg/kg) vs. RS67333 (30 mg/kg) | - | - | 0.0096 | ** | |
| | | | Re-exposure Freezing (%) | RMANOVA | Drug | 5.314 | 3,55 | 0.0027 | ** | 2C |
| | | | | | Time | 11.400 | 4,220 | <0.0001 | **** | |
| | | | | | Drug x Time | 0.798 | 12,220 | 0.6528 | ns | |
| | | | | Fisher's LSD | Saline vs. RS67333 (1.5 mg/kg) | - | - | 0.0185 | * | |
| | | | | | Saline vs. RS67333 (10 mg/kg) | - | - | 0.0006 | *** | |
| | | | | | Saline vs. RS67333 (30 mg/kg) | - | - | 0.2625 | ns | |
| | | | | | RS67333 (1.5 mg/kg) vs. RS67333 (10 mg/kg) | - | - | 0.8278 | ns | |

| | | | | | | | | | | |
|---|---|---|---|---|---|---|---|---|---|---|
| | | | | | RS67333 (1.5 mg/kg) vs. RS67333 (30 mg/kg) | - | - | 0.3259 | ns | |
| | | | | | RS67333 (10 mg/kg) vs. RS67333 (30 mg/kg) | - | - | 0.3050 | ns | |
| | | | Re-exposure Average Freezing (%) | ANOVA | Drug | 5.314 | 3,55 | 0.0027 | ** | 2D |
| | | | | Fisher's LSD | Saline vs. RS67333 (1.5 mg/kg) | - | - | 0.0185 | * | |
| | | | | | Saline vs. RS67333 (10 mg/kg) | - | - | 0.0006 | *** | |
| | | | | | Saline vs. RS67333 (30 mg/kg) | - | - | 0.2625 | ns | |
| | | | | | RS67333 (1.5 mg/kg) vs. RS67333 (10 mg/kg) | - | - | 0.8278 | ns | |
| | | | | | RS67333 (1.5 mg/kg) vs. RS67333 (30 mg/kg) | - | - | 0.3259 | ns | |
| | | | | | RS67333 (10 mg/kg) vs. RS67333 (30 mg/kg) | - | - | 0.3050 | ns | |
| | Forced Swim Test | FST | Day 1 Immobility Time (sec) | RMANOVA | Drug | 3.381 | 3,55 | 0.0245 | * | 2E |
| | | | | | Time | 25.700 | 5,275 | <0.0001 | **** | |
| | | | | | Drug x Time | 0.484 | 15,275 | 0.9479 | ns | |
| | | | | Fisher's LSD | Saline vs. RS67333 (1.5 mg/kg) | - | - | 0.3784 | ns | |
| | | | | | Saline vs. RS67333 (10 mg/kg) | - | - | 0.0028 | ** | |
| | | | | | Saline vs. RS67333 (30 mg/kg) | - | - | 0.8437 | ns | |
| | | | | | RS67333 (1.5 mg/kg) vs. RS67333 (10 mg/kg) | - | - | 0.3430 | ns | |
| | | | | | RS67333 (1.5 mg/kg) vs. RS67333 (30 mg/kg) | - | - | 0.5994 | ns | |
| | | | | | RS67333 (10 mg/kg) vs. RS67333 (30 mg/kg) | - | - | 0.1099 | ns | |
| | | | Day 2 Immobility Time (sec) | RMANOVA | Drug | 0.69 | 3,55 | 0.5620 | ns | 2F |
| | | | | | Time | 5.210 | 5,275 | <0.0001 | **** | |
| | | | | | Drug x Time | 0.720 | 15,275 | 0.7637 | ns | |
| | | | Day 2 Immobility Time (min 3-6) (sec) | ANOVA | Drug | 0.942 | 3,55 | 0.4266 | ns | 2G |
| | Open Field Test | OF | Distance Travelled (cm) | RMANOVA | Drug | 1.368 | 1,18 | 0.2575 | ns | 2H |
| | | | | | Time | 1.513 | 9,162 | 0.1472 | ns | |
| | | | | | Drug x Time | 0.755 | 9,162 | 0.6578 | ns | |
| | | | Time in Center (%) | *t*-test | Saline vs. RS67333 (10 mg/kg) | - | - | 0.2475 | ns | 2I |
| | Elevated Plus Maze | EPM | Time in Open Arms (sec) | *t*-test | Saline vs. RS67333 (10 mg/kg) | - | - | 0.2069 | ns | 2J |
| | | | Entries into Open Arms (sec) | *t*-test | Saline vs. RS67333 (10 mg/kg) | - | - | 0.3584 | ns | 2K |
| | Novelty Suppressed Feeding | NSF | Fraction of mice not eating | Log-rank (Mantel-Cox) test | Drug | - | - | 0.0183 | * | 2L |
| | | | Latency to feed (sec) | *t*-test | Saline vs. RS67333 (10 mg/kg) | - | - | 0.0108 | * | 2M |
| | | | Food Eaten (g) | *t*-test | Saline vs. RS67333 (10 mg/kg) | - | - | 0.3871 | ns | 2N |
| | | | Body Weight Loss (g) | *t*-test | Saline vs. RS67333 (10 mg/kg) | - | - | 0.8829 | ns | 2O |
| | Contextual Fear Conditioning | CFC | Training Freezing (%) | RMANOVA | Drug | 0.7298 | 2,10 | 0.5060 | ns | 3B |
| | | | | | Time | 20.2700 | 4,40 | <0.0001 | **** | |
| | | | | | Drug x Time | 0.835 | 8,40 | 0.5779 | ns | |
| | | | Re-exposure Freezing (%) | RMANOVA | Drug | 0.666 | 2,10 | 0.5351 | ns | 3C |
| | | | | | Time | 7.098 | 4,40 | 0.0002 | ** | |
| | | | | | Drug x Time | 0.665 | 8,40 | 0.7189 | ns | |

| | | | | | | | | | | |
|---|---|---|---|---|---|---|---|---|---|---|
| Figure 3 Acute RS67333 Female | | | Re-exposure Average Freezing (%) | ANOVA | Drug | 0.666 | 2,10 | 0.5351 | ns | 3D |
| | Forced Swim Test | FST | Day 1 Immobility Time (sec) | RMANOVA | Drug | 0.100 | 2,10 | 0.9056 | ns | 3E |
| | | | | | Time | 18.020 | 5,50 | <0.0001 | **** | |
| | | | | | Drug x Time | 1.400 | 10,50 | 0.2076 | ns | |
| | | | Day 2 Immobility Time (sec) | RMANOVA | Drug | 0.185 | 2,10 | 0.8339 | ns | 3F |
| | | | | | Time | 2.952 | 5,50 | 0.0206 | * | |
| | | | | | Drug x Time | 0.556 | 10,50 | 0.8412 | ns | |
| | | | Day 2 Immobility Time (min 3-6) (sec) | ANOVA | Drug | 0.563 | 2,10 | 0.5864 | ns | 3G |

| | | | | | | | | | | |
|---|---|---|---|---|---|---|---|---|---|---|
| | Contextual Fear Conditioning | CFC | Training Freezing (%) | RMANOVA | Drug | 0.317 | 5,34 | 0.8992 | ns | 4B |
| | | | | | Time | 50.650 | 4,136 | <0.0001 | **** | |
| | | | | | Drug x Time | 0.736 | 20,136 | 0.7835 | ns | |
| | | | Re-exposure Freezing (%) | RMANOVA | Drug | 5.284 | 5,34 | 0.0011 | ** | 4C |
| | | | | | Time | 45.700 | 4,136 | <0.0001 | **** | |
| | | | | | Drug x Time | 1.201 | 20,136 | 0.2632 | ns | |
| | | | | Fisher's LSD | Saline vs. (R,S)-ketamine | - | - | 0.0448 | * | |
| | | | | | Saline vs. Prucalopride (3 mg/kg) | - | - | 0.0004 | *** | |
| | | | | | Saline vs. Prucalopride (10 mg/kg) | - | - | 0.4723 | ns | |
| | | | | | Saline vs. PF04995274 (3 mg/kg) | - | - | 0.6871 | ns | |
| | | | | | Saline vs. PF04995274 (10 mg/kg) | - | - | 0.0255 | * | |
| | | | | | (R,S)-ketamine (30 mg/kg) vs. Prucalopride (3 mg/kg) | - | - | 0.1343 | ns | |
| | | | | | (R,S)-ketamine (30 mg/kg) vs. Prucalopride (10 mg/kg) | - | - | 0.1023 | ns | |
| | | | | | (R,S)-ketamine (30 mg/kg) vs. PF04995274 (3 mg/kg) | - | - | 0.1027 | ns | |
| | | | | | (R,S)-ketamine (30 mg/kg) vs. PF04995274 (10 mg/kg) | - | - | 0.8015 | ns | |
| | | | | | Prucalopride (3 mg/kg) vs. Prucalopride (10 mg/kg) | - | - | 0.0004 | *** | |
| | | | | | Prucalopride (3 mg/kg) vs. PF04995274 (3 mg/kg) | - | - | 0.0014 | ** | |
| | | | | | Prucalopride (3 mg/kg) vs. PF04995274 (10 mg/kg) | - | - | 0.2229 | ns | |
| | | | | | Prucalopride (10 mg/kg) vs. PF04995274 (3 mg/kg) | - | - | 0.7981 | ns | |
| | | | | | Prucalopride (10 mg/kg) vs. PF04995274 (10 mg/kg) | - | - | 0.0568 | ns | |
| | | | | | PF04995274 (3 mg/kg) vs. PF04995274 (10 mg/kg) | - | - | 0.0619 | ns | |
| | | | | ANOVA | Drug | 5.284 | 5,34 | 0.0011 | ** | |
| | | | | | Saline vs. (R,S)-ketamine | - | - | 0.0448 | * | |
| | | | | | Saline vs. Prucalopride (3 mg/kg) | - | - | 0.0004 | *** | |
| | | | | | Saline vs. Prucalopride (10 mg/kg) | - | - | 0.4723 | ns | |
| | | | | | Saline vs. PF04995274 (3 mg/kg) | - | - | 0.6871 | ns | |
| | | | | | Saline vs. PF04995274 (10 mg/kg) | - | - | 0.0255 | * | |
| | | | | | (R,S)-ketamine (30 mg/kg) vs. Prucalopride (3 mg/kg) | - | - | 0.1343 | ns | |
| | | | | | (R,S)-ketamine (30 mg/kg) vs. Prucalopride (10 mg/kg) | - | - | 0.1023 | ns | |

| | | | | | | | | | | |
|---|---|---|---|---|---|---|---|---|---|---|
| Figure 4 Acute Prucalopride and PF04995274 Male | | | Re-exposure Average Freezing (%) | Fisher's LSD | (R,S)-ketamine (30 mg/kg) vs. PF04995274 (3 mg/kg) | - | - | 0.1027 | ns | 4D |
| | | | | | (R,S)-ketamine (30 mg/kg) vs. PF04995274 (10 mg/kg) | - | - | 0.8015 | ns | |
| | | | | | Prucalopride (3 mg/kg) vs. Prucalopride (10 mg/kg) | - | - | 0.0004 | *** | |
| | | | | | Prucalopride (3 mg/kg) vs. PF04995274 (3 mg/kg) | - | - | 0.0014 | ** | |
| | | | | | Prucalopride (3 mg/kg) vs. PF04995274 (10 mg/kg) | - | - | 0.2229 | ns | |
| | | | | | Prucalopride (10 mg/kg) vs. PF04995274 (3 mg/kg) | - | - | 0.7981 | ns | |
| | | | | | Prucalopride (10 mg/kg) vs. PF04995274 (10 mg/kg) | - | - | 0.0568 | ns | |
| | | | | | PF04995274 (3 mg/kg) vs. PF04995274 (10 mg/kg) | - | - | 0.0619 | ns | |
| | Forced Swim Test | FST | Day 1 Immobility Time (sec) | RMANOVA | Drug | 0.520 | 5,34 | 0.7596 | ns | 4E |
| | | | | | Time | 19.370 | 5,170 | <0.0001 | **** | |
| | | | | | Drug x Time | 0.990 | 25,170 | 0.4827 | ns | |
| | | | Day 2 Immobility Time (sec) | RMANOVA | Drug | 3.135 | 5,34 | 0.0197 | * | 4F |
| | | | | | Time | 3.161 | 5,170 | 0.0094 | ** | |
| | | | | | Drug x Time | 0.859 | 25,170 | 0.6616 | ns | |
| | | | | Fisher's LSD | Saline vs. (R,S)-ketamine (30 mg/kg) | - | - | 0.0218 | * | |
| | | | | | Saline vs. Prucalopride (3 mg/kg) | - | - | 0.0052 | ** | |
| | | | | | Saline vs. Prucalopride (10 mg/kg) | - | - | 0.1772 | ns | |
| | | | | | Saline vs. PF04995274 (3 mg/kg) | - | - | 0.1259 | ns | |
| | | | | | Saline vs. PF04995274 (10 mg/kg) | - | - | 0.0023 | ** | |
| | | | | | (R,S)-ketamine (30 mg/kg) vs. Prucalopride (3 mg/kg) | - | - | 0.8332 | ns | |
| | | | | | (R,S)-ketamine (30 mg/kg) vs. Prucalopride (10 mg/kg) | - | - | 0.1712 | ns | |
| | | | | | (R,S)-ketamine (30 mg/kg) vs. PF04995274 (3 mg/kg) | - | - | 0.4094 | ns | |
| | | | | | (R,S)-ketamine (30 mg/kg) vs. PF04995274 (10 mg/kg) | - | - | 0.3813 | ns | |
| | | | | | Prucalopride (3 mg/kg) vs. Prucalopride (10 mg/kg) | - | - | 0.0568 | ns | |
| | | | | | Prucalopride (3 mg/kg) vs. PF04995274 (3 mg/kg) | - | - | 0.2475 | ns | |
| | | | | | Prucalopride (3 mg/kg) vs. PF04995274 (10 mg/kg) | - | - | 0.4224 | ns | |
| | | | | | Prucalopride (10 mg/kg) vs. PF04995274 (3 mg/kg) | - | - | 0.6674 | ns | |
| | | | | | Prucalopride (10 mg/kg) vs. PF04995274 (10 mg/kg) | - | - | 0.0209 | * | |
| | | | | | PF04995274 (3 mg/kg) vs. PF04995274 (10 mg/kg) | - | - | 0.0941 | ns | |
| | | | | ANOVA | Drug | 2.940 | 5,34 | 0.0260 | * | |
| | | | | | Saline vs. (R,S)-ketamine (30 mg/kg) | - | - | 0.0330 | * | |
| | | | | | Saline vs. Prucalopride (3 mg/kg) | - | - | 0.0093 | ** | |
| | | | | | Saline vs. Prucalopride (10 mg/kg) | - | - | 0.2443 | ns | |
| | | | | | Saline vs. PF04995274 (3 mg/kg) | - | - | 0.1050 | ns | |

| | | | | | | | | | | |
|---|---|---|---|---|---|---|---|---|---|---|
| | | | Day 2 Immobility Time (min 3-6) (sec) | Fisher's LSD | Saline vs. PF04995274 (10 mg/kg) | - | - | 0.0027 | ** | 4G |
| | | | | | (R,S)-ketamine (30 mg/kg) vs. Prucalopride (3 mg/kg) | - | - | 0.8478 | ns | |
| | | | | | (R,S)-ketamine (30 mg/kg) vs. Prucalopride (10 mg/kg) | - | - | 0.1763 | ns | |
| | | | | | (R,S)-ketamine (30 mg/kg) vs. PF04995274 (3 mg/kg) | - | - | 0.5815 | ns | |
| | | | | | (R,S)-ketamine (30 mg/kg) vs. PF04995274 (10 mg/kg) | - | - | 0.3207 | ns | |
| | | | | | Prucalopride (3 mg/kg) vs. Prucalopride (10 mg/kg) | - | - | 0.0622 | ns | |
| | | | | | Prucalopride (3 mg/kg) vs. PF04995274 (3 mg/kg) | - | - | 0.4089 | ns | |
| | | | | | Prucalopride (3 mg/kg) vs. PF04995274 (10 mg/kg) | - | - | 0.3387 | ns | |
| | | | | | Prucalopride (10 mg/kg) vs. PF04995274 (3 mg/kg) | - | - | 0.4653 | ns | |
| | | | | | Prucalopride (10 mg/kg) vs. PF04995274 (10 mg/kg) | - | - | 0.0156 | * | |
| | | | | | PF04995274 (3 mg/kg) vs. PF04995274 (10 mg/kg) | - | - | 0.1270 | ns | |
| | Open Field | OF | Distance Traveled (cm) | RMANOVA | Drug | 0.350 | 45,306 | 0.9139 | ns | 4H |
| | | | | | Time | 7.441 | 9,306 | <0.0001 | **** | |
| | | | | | Drug x Time | 0.7151 | 45,306 | 0.9139 | ns | |
| | Elevated Plus Maze | EPM | Time in Open Arms (sec) | ANOVA | Drug | 1.870 | 5,34 | 0.1257 | ns | 4I |
| | | | Entries Into Open Arms (sec) | ANOVA | Drug | 1.599 | 5,34 | 0.1686 | ns | 4J |
| | Novelty Suppressed Feeding | NSF | Fraction of mice not eating | Log-rank (Mantel-Cox) test | Drug | - | - | 0.0316 | * | 4K |
| | | | Latency to Feed (sec) | ANOVA | Drug | 2.115 | 5,34 | 0.0874 | ns | 4L |
| | | | Body Weight Loss (g) | ANOVA | Drug | 1.293 | 5,34 | 0.2899 | ns | 4M |
